# Genomic and phylogenetic analysis of the first myovirus isolated from *Oceanospirillaceae*, representing a novel viral cluster prevalent in polar oceans

**DOI:** 10.1101/2022.01.27.478124

**Authors:** Wenjing Zhang, Yundan Liu, Jinyan Xing, Kaiyang Zheng, Qian Li, Chengxiang Gu, Ziyue Wang, Hongbing Shao, Cui Guo, Hui He, Hualong Wang, Yeong Yik Sung, Wen Jye Mok, Li Lian Wong, Yantao Liang, Andrew McMinn, Min Wang

**Author notes:** Correspondence (Y.L.); (M.W.). These authors contributed equally to this work (Wenjing Zhang, Yundan Liu, Jinyan Xing).

## Abstract

The marine bacterial family *Oceanospirillaceae*, which is abundant in the deep-seas and polar oceans, is closely associated with algal blooms and petroleum hydrocarbons degradation. However, only a few *Oceanospirillaceae*-infecting phages have so far been reported. Here we report on a novel *Oceanospirillum* phage, vB_OsaM_PD0307, which is the first myovirus to be found that infects *Oceanospirillaceae*. vB_OsaM_PD0307 with a 44,421 bp linear dsDNA genome. Phylogenetic analysis and average nucleotide sequence identities suggest that vB_OsaM_PD0307 is different from other phage isolates and represents a novel genus-level myoviral cluster with two high-quality uncultured viral genomes, designed as *Oceanospimyovirus*. Additionally, the biogeographical distribution of the vB_OsaM_PD0307 cluster suggests that they are widespread in the oceans and abundant in polar areas. In summary, our findings expand the current understanding of the phylogenetic diversity, genomic characteristic and function of *Oceanospimyovirus* phages, and highlight the role of the vB_OsaM_PD0307 phage as a major ecological agent that can infect certain key bacterial groups associated with polar algal blooms.

**Importance:** Oceanospirillumphage vB_OsaM_PD0307 is the first myovirus found to infect *Oceanospirillaceae* and represents a novel viral genus, *Oceanospimyovirus*. This study provides insights into the genomic, phylogenetic, and ecological characteristics of myoviruses infecting *Oceanospirillaceae* and improves our understanding of the interactions between *Oceanospirillaceae* and their phages in the oceans.

## Introduction

Viruses are the most abundant and diverse “life forms” in the ocean (1, 2). They mediate fluctuations in microbial abundance and shape community structure and aggregation through the lysis of their hosts, processes referred to as the “viral shunt” and “viral shuttle” (3–5). Heterotrophic bacteria-infecting phage in particular, can cause over 50% mortality rate of their hosts through cell lysis, and this number can be higher in certain environments (6). Viral-host interactions promote marine biogeochemical cycles and marine carbon sequestration through the processes known as the “biological pump” and “microbial carbon pump”. Phages also promote the active evolution and shape phylogeny of their hosts through horizontal gene transfer (7, 8). Over the past few decades, our understanding of viral diversity has significantly expanded, though mostly relying on the development of high-throughput sequencing and metagenomic technology. As a consequence, over 90% of available viral genomes remain uncultured and uncharacterized (there in after referred to as UViGs) (9). Targeted phage isolation methods are vital in terms of filling the knowledge gaps between sequences information and functionality, especially in the context of interaction dynamics and co-evolution between phage and host, as well as shedding light on their potential role in marine microbial food webs..

The bacteria genus *Oceanospirillum*, Hylemon was separated from the genus *Spirillum* based on their differing physiological properties and DNA base composition. The mean GC content of *Oceanospirillum* genomes ranges from 42 to 51 mol% (10, 11). Currently, twenty-three members have been validly published (https://www.bacterio.net/genus/Oceanospirillum) and these have been isolated from a range of marine environments, including coastal waters (10, 12), decaying seaweed (13), mangrove sediments (14, 15), putrid infusions of marine mussels (11, 16, 17) and the leaves of the seagrass (18). Bacteria within this family are well known for their capability of hydrocarbon degradation (19–22) and so are often detected at high abundances in oil-contaminated marine environments (23–25). Recently, a high abundance of *Oceanospirillaceae* was detected in the hadal zone of the Mariana Trench (21), suggesting their potential importance in these extreme marine habitats. In addition, *Oceanospirillaceae* are also prevalent in polar oceans, such as the Amundsen Sea polynya (26, 27), Ross Sea (28) and coastal waters of the Arctic (29). Members of *Oceanospirillaceae* possess a complete B12 vitamin synthesis pathway, which affects DMSP synthesis and promotes algal growth (30), suggesting that *Oceanospirillaceae* plays an important role in some algal blooms. Despite the ecological significance of *Oceanospirillaceae* in the ocean, our understanding of their co-occurring phages is still poor. Currently, only eight phages infecting *Oceanospirillaceae* have been isolated, including three autographiviruses infecting *Marinomonas*, two siphoviruses, and one corticovirus, and two unclassified phages (*Marinomonas* phage MfV and *Nitrincola* phage 1M3-16). There have been no previous reports of Myovirus infecting *Oceanospirillaceae*.

In this study, the first myovirus infecting *Oceanospirillaceae*, named vB_OsaM_PD0307 was isolated and characterized its genomic and phylogenetic features. Phylogenetic analysis showed that vB_OsaM_PD0307 was distantly related to other reported viral isolates in the NCBI dataset and may represent a novel myoviral genus with two high-quality UViGs. The biogeographic distribution analysis suggests that the homologous of vB_OsaM_PD0307 are abundant in polar oceans. This study provides an insight into the genome of *Oceanospirillaceae* myovirus, shedding light on the important interactions between algal bloom-associated bacteria and myoviruses in polar oceans.

## Materials and methods

### Isolation and purification of host phage strain

Both *Oceanospirillum* sp. PD0307 and its phage vB_OsaM_PD0307 were isolated from a surface water sample collected from coastal waters of the Yellow Sea (120°19’32.6”E, 36°4’1.7”N) in June 2020. The host strain was isolated from and maintained in 2216E medium (peptone 5 wt.%, yeast extract 1 wt.%), at 28 ^°^C and 120 rpm, in a shaking incubator.

For phage isolation, 10 ml seawater sample collected from the same station was passed through a 0.22 µm membrane filter to remove any cellular organisms (Isopore™ 0.2 µm GTTP; Merck, Ireland) (31). Phage vB_OsaM_PD0307 was isolated by plaque assay using the double-layer plating method. Briefly, 200 μl seawater filtered 0.22 µm pore-size filters was mixed with the host culture (approximately 8-hour) and incubated for 30-minute, allowing the absorption of the phages at room temperature. Then, 4 ml of the semi-solid culture (at 55 ^°^C) was added to the mixture, pouring it onto the plate after vortexing. Plates were cultivated at 28 ^°^C and monitored until visible plaques were formed in the double layer culture (usually happens within 24 h).

A single plaque was picked from the double plate and suspended in 2 ml of SM buffer, (100 mM of NaCl, 8 mM of MgSO4, 50 mM of Tris-HCl, at pH 7.5) and then purified three times via plaque assay. Culture lysates were recovered to enrich and concentrate the phages. Approximately 500 ml of exponentially growing host were challenged with 50ml purified viral stock and incubated at 28 ^°^C for 24 h. The lysate was first filtered through a 0.22 µm membrane filter to remove uninfected host cells, then concentrated from 500 ml to 5 ml using 30 kDa super-filters (UFC5030, Millipore). Concentrated phage lysate was treated with a second filtration by passing through 0.22 µm Supor membrane.

### Preparation for phage morphology observation

Phage particles were precipitated by adding PEG 8000 and NaCl to a final concentration of 10% (w/v) and 1 M, respectively, and incubated overnight at 4°C. Phage particles were precipitated by centrifugation at 15,000 g for 30 min and then re-suspended in 5 ml of SM buffer, 20 μl of the mixture was then taken and placed on a copper net and stained with 2 wt.% phosphotungstic acids (pH 7.5) for 5 min. Its morphology was identified using transmission electron microscope (TEM) (JEOLJEM-1200EX, Japan) at 100 KV, equipped with a diamond knife for thin sectioning, and Images were taken using GATAN INC CCD image transmission system (Gatan Inc., Pleasanton, CA, USA).

### Phage DNA preparation, genome sequencing and gene annotation

The concentrated phage lysate was prepared for phage genomic DNA extraction was performed by Virus DNA Kit (OMEGA), according to the manufacturer’s instructions, and quality control was subsequently carried out on the purified DNA samples. The high-quality DNA sample (OD260/280=1.8∼2.0, >6ug) was used to construct the fragment library and then used for Illumina NovaSeq 6000 sequencing by Shanghai Biozeron Biotechnology Co., Ltd. (Shanghai, China.). The raw paired-end reads were trimmed and quality controlled by Trimmomatic (v. 0.3.6) with parameters: SLIDINGWINDOW:4:15, MINLEN:75 (32). ABySS was used to assemble the viral genome after the quality control processes, multiple-Kmer parameters were chosen to obtain the optimal assembly results (33). GapCloser software was subsequently applied to fill in the remaining local inner gaps and to correct the single base polymorphism for the final assembly and further analysis (34).

Gene models were identified using GeneMarkS (35). Then all gene models were blastp against non-redundant (NR) in the NCBI database, SwissProt (http://uniprot.org), KEGG (http://www.genome.jp/kegg/), and COG (http://www.ncbi.nlm.nih.gov/COG) for functional annotation by the blastp module. The genome visualization was conducted by CLC Main Workbench (v6.8 downloaded on http://www.clcbio.com). In addition, tRNA was identified using the tRNAscan-SE (v1.23) (36) and rRNA was determined using RNAmmer (v1.2 downloaded on https://services.healthtech.dtu.dk/service.php/RNAmmer-1.2). GC skew analysis was performed on Genskew (https://genskew.csb.univie.ac.at/webskew). Average amino acid identity (AAI) between vB_OsaM_PD0307 and other viral sequences was calculated by the AAI calculator (http://enve-omics.ce.gatech.edu) to estimate the distribution of AAI between proteins from two genomic sequences.

### Phylogenic tree construction of host *Oceanospirillaceae* sp

A total of 355 16S rRNA sequences of *Oceanospirillaceae* with defined taxonomy in GenBank, including the host strain Oceanospirillum sp. PD0307, were retrieved from GenBank and aligned by MAFFT (37) using G-INS-1 of strategy with 1000 iterations (mafft --globalpair --maxiterate 1000 16S_Oceanospirillaceae_pro.fasta > 16S_Oceanospirillaceae.mafft). The likelihood phylogenic tree was calculated from multiple sequence alignments using IQ-tree2 (38), applying the GTR+I+G model with 1000 bootstrap iterations (Command: iqtree -s 16S_Oceanospirillaceae.mafft -m MFP -B 1000 -T AUTO) and visualized by iTOL v4 (39).

### Homologous sequence recruitment of phage vB_OsaM_PD0307

A total of 57212 isolated complete phages genomes were downloaded from NCBI GenBank to build a reference isolated-phages dataset (https://www.ncbi.nlm.nih.gov/labs/virus). The whole genome sequences of phage vB_OsaM_PD0307 as a input to find the other similar genomes by blastn in this reference isolated-phages dataset with e-value < 1e-5, identity > 50%. At the same time, homology recruitment in uncultured virus databases was also taking place. The whole genome sequence of phage vB_OsaM_PD0307 was queried against the IMG/VR v3 (40) database using blastn to search for homologous contig sequences (threshold: e-value < 1e-5, percentage of identity > 70%) and seven UViGs were retrieved.

All the isolated and uncultured homologous sequences with vB_OsaM_PD0307, and all eight isolated *Oceanospirillaceae* phages were combined to calculate intergenomic similarity by VIRIDIC (41), as well as ANI via OAT software using the orthogonal method to determine the overall similarity (42). AAI was calculated on the website (http://enve-omics.ce.gatech.edu), which estimated the distribution of AAI between proteins from the two genomic sequences.

### Phylogenetic and comparative genomic analysis

A proteomic tree based on the similarities of the whole genome was generated using VIPTree (43). Each encoding nucleic sequence as a query was searched against the Virus-Host DB using tBLASTx. All viral sequences in Virus-Host DB were selected to generate a first circular tree. The second more accurate phylogenetic tree was regenerated using 35 related phages automatically selected from the first results. 20 isolated sequences were selected as references, and seven *Mycobacterium* phages appeared as an outgroup, vB_OsaM_PD0307 and the other seven UViGs were used as queries to construct the whole-genome phylogenetic tree using VIPTree (43). The group of seven UViGs, *Shewanella* phage SppYZU01, and vB_OsaM_PD0307 were selected to perform the multi-genomic alignments.

### Global oceanic distribution of phage vB_OsaM_PD0307 and relative viral sequences

Stations in Global Ocean Viromes 2.0 (GOV 2.0) were divided into five viral ecological zones (VEZs), including the Arctic (ARC), Antarctic (ANT), temperate and tropical epipelagic (EPI), temperate and tropical mesopelagic (MES), and bathypelagic (BATHY). Five representative stations were selected for each VEZs to assess the relative abundance of vB_OsaM_PD0307 and its relative viral sequences. (ANT: ERR594377, ERR594409, ERR599352, ERR599364, ERR599384; ARC: ERR2762158, ERR2762161, ERR2762163, ERR2762165, ERR2762169; EPI: ERR594353, ERR594398, ERR594395, ERR594403, ERR594376; MES: ERR2752153, ERR2752154, ERR2752163, ERR599375, ERR599379; BATHY: msp112, msp121, msp131, msp144, msp81) The global oceanic distribution was calculated by the metagenomics tool minimap2 (parameters: -min-read-percent-identity 0.95, -min-read-aligned-percent 0.75, -m rpkm) (46) and expressed by RPKM (reads per kilobase per million mapped reads) values. Besides, pelagiphage HTVC010P, HTVC011P, cyanophages P-SSP7, P-SSM7, S-SM2, S-CBS2, roseophage SIO1, SAR116 phage HMO2011 which have significantly representative in different oceanic areas and depths as the references.

### Data availability

The complete genome of *Oceanospirillum* phage vB_OsaM_PD0307 has been deposited in NCBI GeneBank under accession number OL658619, and the 16S rRNA sequence of the host also has been deposited in NCBI GeneBank under accession number OL636378.

## Results and Discussion

### Isolation and morphology of the plaques of vB_OsaM_PD0307

The phage vB_OsaM_PD0307 (accession: OL658619), infecting *Oceanospirillum* sp. PD0307 (accession: OL636378), was isolated from a surface seawater sample from the Yellow Sea. The morphology of vB_OsaM_PD0307 was that of a myovirus with an icosahedral head of 51 ± 2 nm in length and a 112 ± 3 nm-long contractile tail (Fig. 1A). Infection of vB_OsaM_PD0307 formed clear and round (1–2 mm diameter average) plaques in double-layer culture (Fig. 1B).

**Fig. 1.**
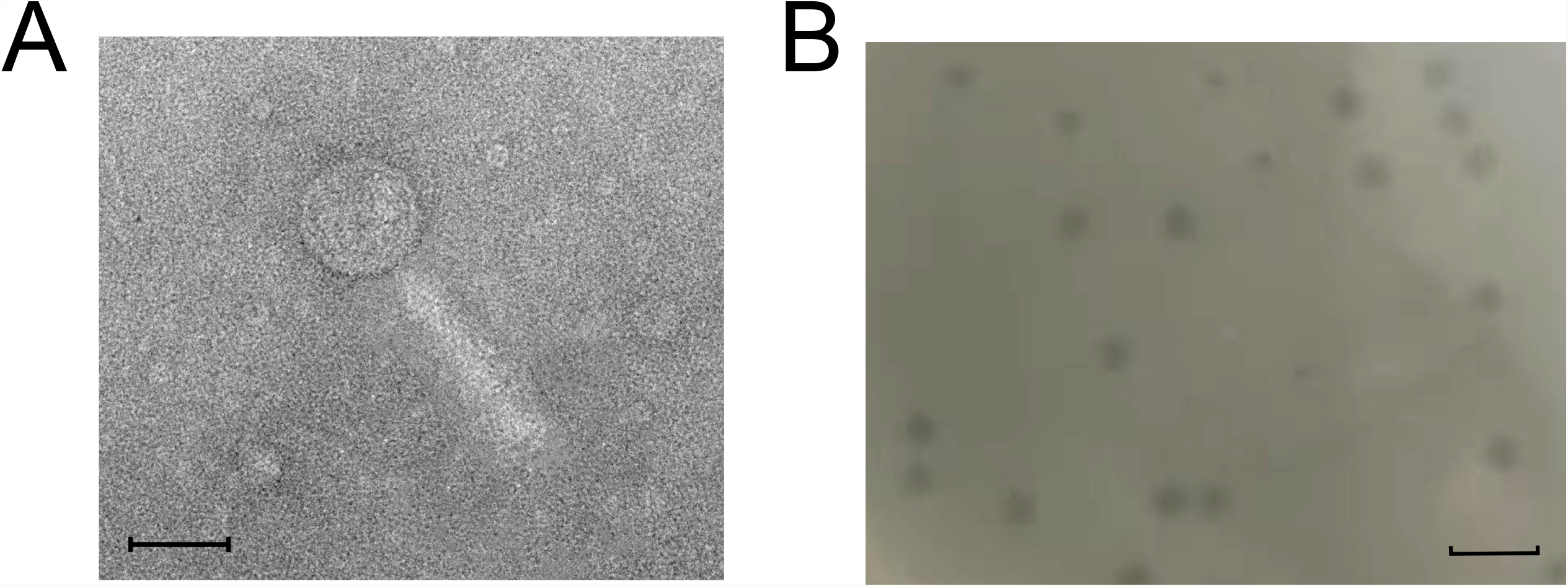
(A) Morphology and biological properties of phage vB_OsaM_PD0307, the scale bar is 50 nm. (B) Phage plaques of vB_OsaM_PD0307 formed in double-layer agar plates, the scale bar is 5 mm

### Phylogenetic analysis of the host bacterium *Oceanospirillum* sp. PD0307

The phylogenic position of the host bacterium *Oceanospirillum* sp. PD0307 was clustered with other *Oceanospirillaceae* strains in the phylogenetic tree. The monophyletic clade containing *Oceanospirillum* sp. PD0307 includes four genera (*Neptuniibacter, Profundimonas, Amphritea*, and *Oceanospirillum*), indicating its close relationship with other *Oceanospirillaceae*, especially with other *Oceanospirillum* strains (Fig. 2). Although *Oceanospirillum* sp. PD0307 is on a sister branch with *Oceanospirillum sanctuarii* AK56, which was isolated from sediment (14), the branch with *Oceanospirillum* sp. PD0307 has a relatively distant phylogenic link with them, suggesting *Oceanospirillum* sp. PD0307 might be a novel species of *Oceanospirillum*. In addition, *Oceanospirillum* sp. PD0307 has the longest branch length in the tree (0.121), indicating that *Oceanospirillum* sp. PD0307 might have evolved from a common ancestor of *Oceanospirillum* or even *Oceanospirillaceae*. Thus, the interaction between phage vB_OsaM_PD0307 and *Oceanospirillum* sp. PD0307 might represent a novel case in the co-evolutionary history between viruses and *Oceanospirillum*.

**Fig. 2.**
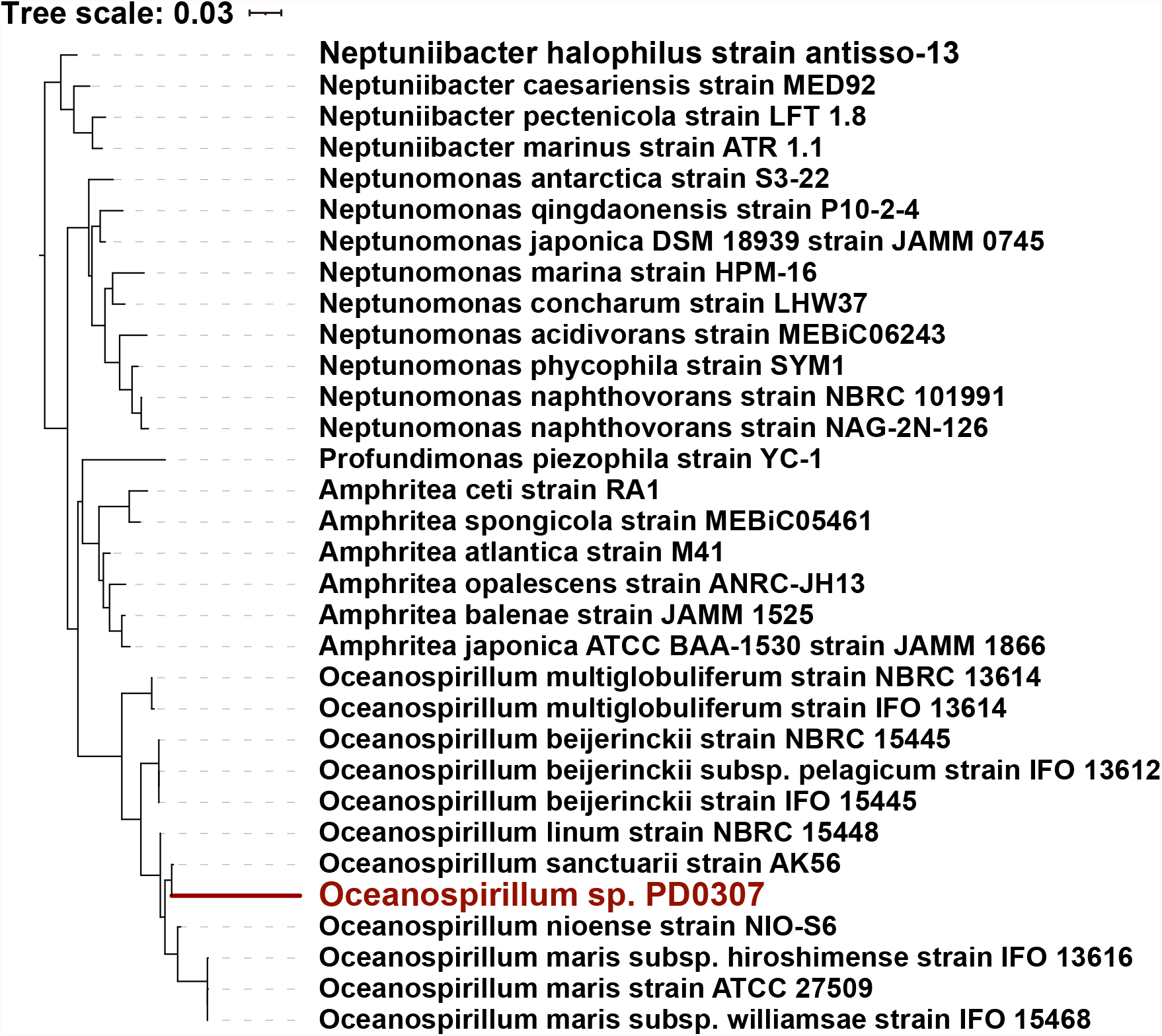
The maximum likelihood phylogenic tree of *Oceanospirillum* sp. PD0307 and 16S rRNA sequences of 31 related *Oceanospirillaceae* species. The *Oceanospirillum* sp. PD0307 was highlighted in the tree. A relatively close relationship of *Oceanospirillum* sp. PD0307 with other *Oceanospirillum* was displayed, while monophyly of its branch was observed.

### Genomic features of phage vB_OsaM_PD0307

According to the genomic sequencing and assembly results, phage vB_OsaM_PD0307 has a 44,421-bp linear dsDNA genome with a GC content of 57.13%. No tRNA and rRNA genes were found in the genome. The genome had a 96.41% encoding rate consisting of 56 predicted open reading frames (ORFs). Thirty-seven ORFs are located on the sense strand, accounting for 66.07% of the total coding genes, and nineteen ORFs on the antisense strand (Fig. 3A, Table S1). There were 11 coding regions (19.64 % ORF angenes) that did not match any homologous sequence under the restriction of *e*-value < 1e-5 in all 56 genes. Among the remaining 45 genes that matched homologous sequences, 23 identified specific functions, and 22 matched homologous proteins containing unknown functions. The 23 functional ORFs could be classified into three different modules: twelve ORFs for DNA replication and metabolism, ten ORFs for phage structure and packing proteins, and one auxiliary metabolic gene (AMG, transcriptional regulator) (Fig. 3A).

**Fig. 3.**
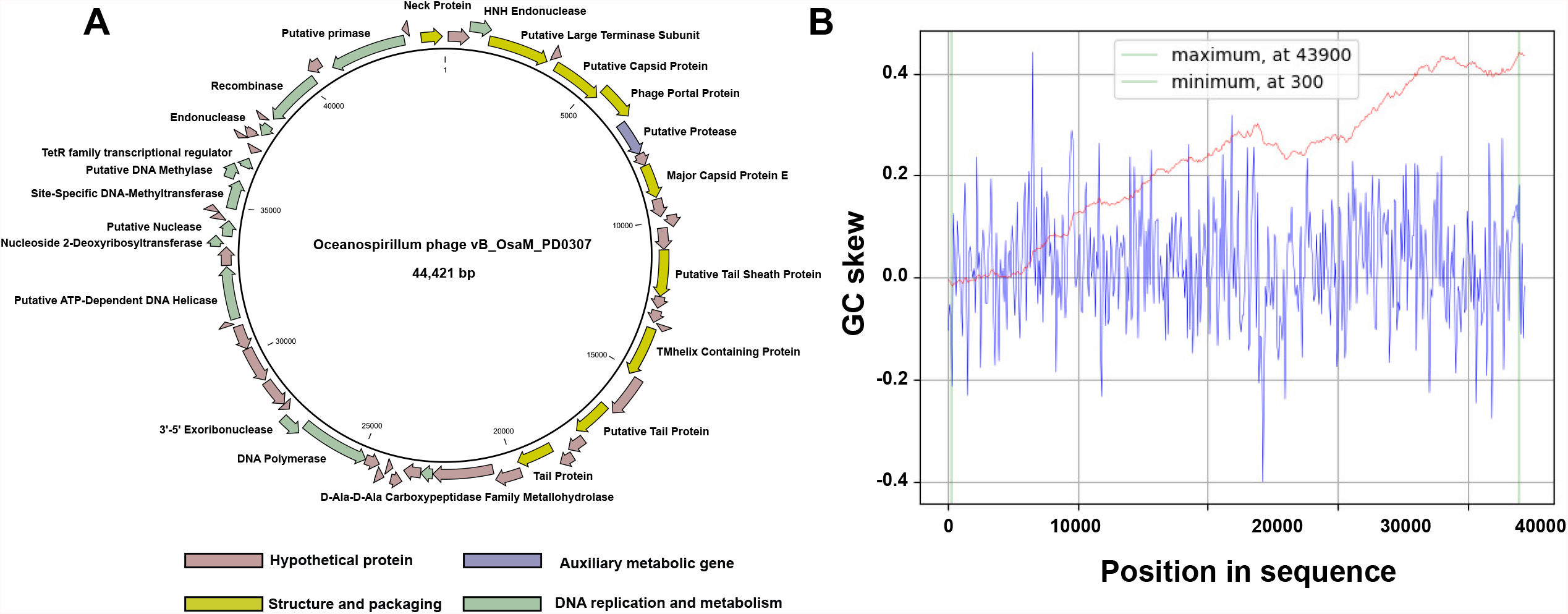
Circularized genome map (A) and cumulative GC skew analysis (B) of the genome sequence of Oceanospirillumphage vB_OsaM_PD0307. (A) The outer circle represents different categories of putative functional genes, which were represented by different colors. (B) the cumulative graph of minimum and maximum values of GC skew were displayed and calculated by using a window size of 1,00 bp and a step size of 100 bp. The GC-skew and the cumulative GC-skew were represented by blue and red lines, respectively. The minimum and maximum of a GC-skew could be used to predict the origin of replication (300 nt) and the terminus location (43900 nt).

In the genome of phage vB_OsaM_PD0307, twelve ORFs were predicted to encode genes related to DNA replication and metabolism. ORF 25 encoded the D-Ala-D-Ala carboxypeptidase family metallohydrolase gene, and also showed high homology with hedgehog signaling/DD-peptidase zinc-binding domain gene of *Vibrio* phage 1.169.O._10N.261.52.B (pident 58.6, qcovhsp 100, bitscore 154.1, evalue 4.4E-34). The structure of the N-terminal signaling domain of hedgehog proteins has been resolved and reveals a tetrahedrally coordinated zinc ion, which is structurally homologous to the zinc-binding motif in bacterial D-alanyl-D-alanine carboxypeptidases (DD-peptidases) (44–46). ORF 25 contained some amino acids of peptidase genes, commonly detected within some bacterial genomes, such as motifs HXXXXXXD and WXH, which were typical motifs for peptidase M15 subfamily A (47, 48). ORFs 44 and 45 encoded DNA-methyltransferase and DNA methylase genes, respectively. Site-specific DNA-methyltransferase, N-6 adenine-specific DNA methylase, and cytosine-N4-specific are enzymes that specifically methylate the amino group at the C-4 position of cytosines and the N-6 position of adenine in DNA. They utilize the cofactor S-adenosyl-L-methionine as the methyl donor and are active as monomeric enzymes. In prokaryotes, the major role of DNA methylation is to protect host DNA against degradation by restriction enzymes (49, 50), and DNA methylation of phage vB_OsaM_PD0307 may have a similar function to elude host immunity.

Ten ORFs encoding genes related to the structure and packaging modules are located at the front end of the genome. The putative protease gene (ORF 7) is affiliated with the cl23717 superfamily, which has portal proteins upstream and capsid proteins downstream. Capsid maturation in double-stranded-DNA (dsDNA) phages requires proteolytic cleavage by a prohead protease (51). In the BLAST results, ClpP/crotonase-like domain proteins were significantly matched to the S49 family proteins, members of the large crotonase superfamily (52). One of the typical features of myotail phages is the presence of tail sheath proteins. ORF 13 encoded these in Phage vB_OsaM_PD0307, which had a strong homologous sequence with the myophage *Shewanella* phage SppYZU01 with 99% qcov and 66.87% pident. The top 50 homologous sequences are all present in the myophages under the given thresholds (*e*-value < 1e-50, qcov > 90%, pident > 30%). The TMhelix (ORF 17) containing gene, which is sandwiched between two tail genes, is related to the transport of substances across cell membranes (53) and may be related to the adsorption of host bacteria by phages.

Only one AMG was detected in the genome of phage vB_OsaM_PD0307 and this encoded a gene related to the TetR family transcriptional regulator (ORF 46). It is well represented and widely distributed among bacteria with an HTH DNA-binding motif (54). Members of this family are well known for their roles as regulators of antibiotic efflux pumps. This gene is a phage-mediated transcriptional regulator for antimicrobial resistance and can help host cells to survive in antimicrobial environments (55).

Cumulative GC skew analyses were performed to determine the origin and terminus of replication of the phage genome (56). The minimum GC skew was at 300 nt and the maximum at 43900 nt, which are at the head and tail of the genome respectively (Fig. 3B). Two inflection points were identified in the above regions, indicating an asymmetric base composition, which were lowest at the origin and the highest at the terminus.

### Phylogenetic and synteny analysis between phage vB_OsaM_PD0307 and its homologous sequences

To further understand the phylogenetic relationship between phage vB_OsaM_PD0307 and other isolated phages, 1,812 dsDNA phage genomes were selected from the Virus-Host database to establish the whole-genome phylogenetic tree. Of these, phage vB_OsaM_PD0307 originated from the tree root and formed a separate clade (Fig. 4A). The detailed phylogenetic trees were then regenerated after adding the seven homologous UViGs. Phage vB_OsaM_PD0307, *Shewanella* phage SppYZU01, and the seven homologous UViGs were grouped together and formed a unique viral cluster (Figs. 4B and 4C). Among them, S85_DCM_NO_526, S137 and UViG_281 were very close to vB_OsaM_PD0307 and shared the same ancestral branch, indicating that they had the same evolutionary characteristics and similar genetic relationships.

**Fig. 4.**
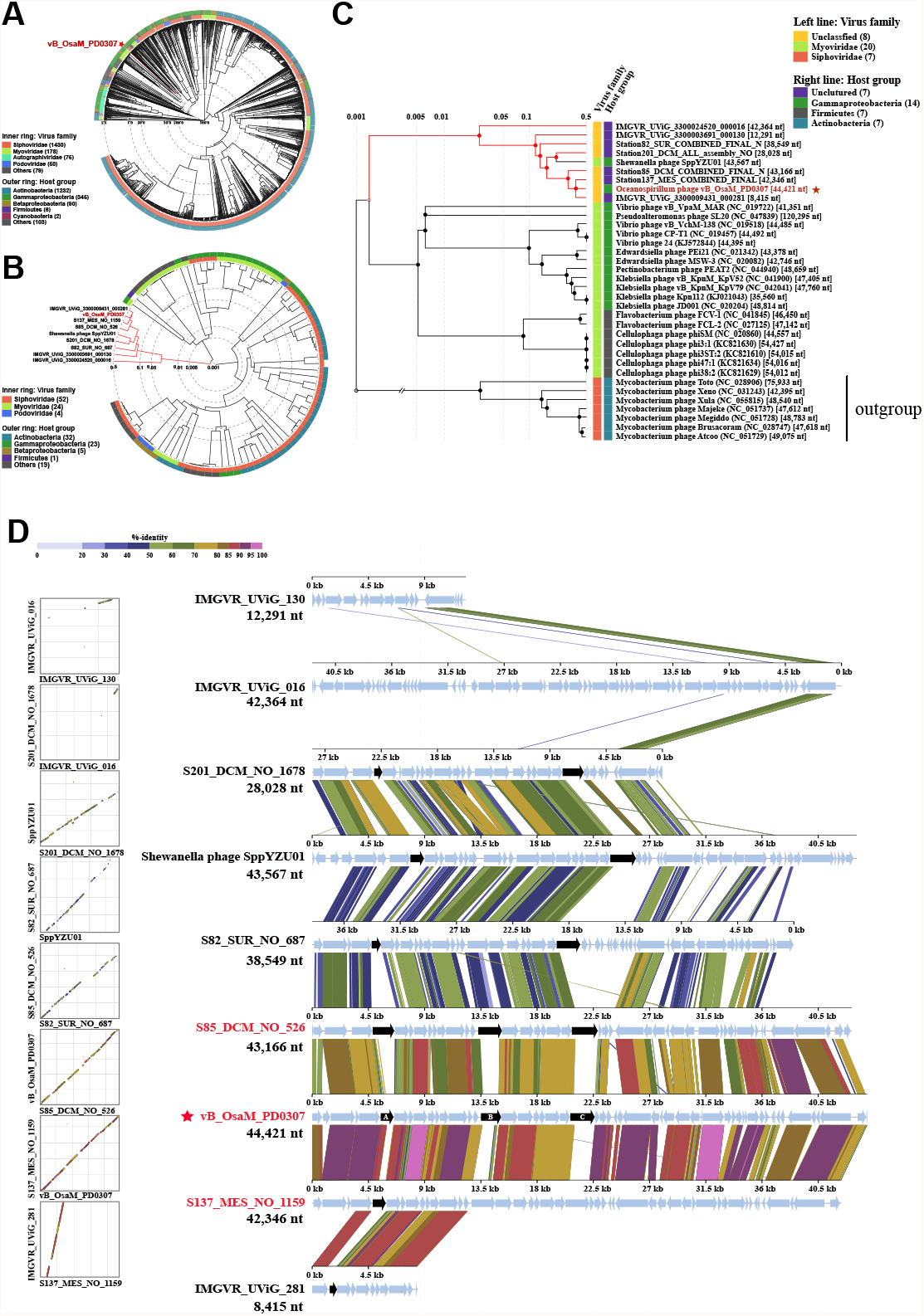
Phylogenetic trees of Oceanospirillum Phage vB_OsaM_PD0307 with different references. (A) The whole-genomes phylogenetic tree was constructed with 1,812 dsDNA phages genomes in the Virus-Host database as references. (B) Nine *Oceanospimyovirus* (Oceanospirillum Phage vB_OsaM_PD0307, Shewanella phage SppYZU01, and seven homologous uncultured viral genomes) as queries and other 80 related phages were used to construct a circular phylogenetic tree. (C) A rectangular phylogenetic tree was established with *Oceanospimyovirus* and seven Mycobacterium phages, which were used as outgroups for control. (D) Gene synteny of Oceanospirillum Phage vB_OsaM_PD0307, Shewanella phage SppYZU01, and seven homologous uncultured viral genomes in IMG/VR v3 database. Sequences comparison performed using tBLASTx (10 bp minimum alignment) with percent identity. Synteny was recognized when genomes featured a minimum of five consecutive syntenic genes within the same genomic area and separated by a maximum of four non-syntenic genes.

To clarity the common features of phage vB_OsaM_PD0307 and homologous viral genomes, an alignment of these sequences by tBLASTx was performed. Comparative genomic analysis revealed that they had a universal homology at the amino acid level, especially among vB_OsaM_PD0307, S85_DNC_NO_526 and S137_MES_NO_1159 (Fig. 4D). These three viral genomes showed a high degree of synteny in the whole-genome arrangement, indicating correlations in the phylogenetic process. Most structural and packaging gene similarities suggest that these viruses may be taxonomically similar. Given that this is the first isolate of this group of viruses, it is likely that this is a new and undiscovered group of viruses. Even the only AMG of vB_OsaM_PD0307, the TetR family transcriptional regulator gene, can be found in a similar region to the other two homologous viral genomes. This suggests that this viral group may possess the ability to assist host cells survive environmental antibiotics. It is worth noting that genes A, B and C of vB_OsaM_PD0307, which encode the portal protein, TMhelix containing protein and hypothetical protein respectively, were common but not homologous among the group; this may relate to the range of hosts of these viruses.

### vB_OsaM_PD0307 represented a new myoviral genus

Based on the results of homology search in the reference isolated-phages dataset, only one genome, Shewanella phage SppYZU01, was retrieved, with a relatively low average amino acid identity (AAI, 48.33%) and average nucleotide sequence identity (ANI, 62.16%). Therefore, BLASTn was used to search the IMG/VR v3 (40) database to find the homologous UViG sequences. In total seven UVIGs, under the thresholds (min pident 70.21%, and min e-value 1.60E-08), were screened. They were all assembled from marine waters or sediment samples, three of them were judged as high-quality sequences (i.e. genome ≥90% complete and non-redundant) and four contigs were judged as genome fragments (i.e. genome <90% complete). The longest and shortest sequences were 43.16 and 8.41 kb, respectively (Table S2).

All eight complete genomes of the isolated *Oceanospirillaceae*-infecting phages and the eight homologous sequences with vB_OsaM_PD0307 were combined and the intergenomic similarity of the seventeen sequences were analysed using VIRIDIC (41). Three *Marinomonas* phages assigned to the same genus, *Murciavirus* shared high intergenomic similarity (> 95%), and their aligned genome fraction and genome lengths were all 100%. In addition, the viral genomes infecting *Oceanospirillaceae* are diverse and almost all were quite different from each other (Fig. 5). Of the eight vB_OsaM_PD0307 related homologous UViGs, three sequences in the red box of the heatmap shared a high intergenomic similarity (> 50%) and had a similar aligned genome fractions and genome lengths (Fig. 5A). IMGVR_UViG_281 and vB_OsaM_PD0307 had a higher aligned genome fraction (80%), but their intergenomic similarity was low because of the differences in their genome length. The genome length of IMGVR_UViG_281 was only 8,415bp (Table S2).

**Fig. 5.**
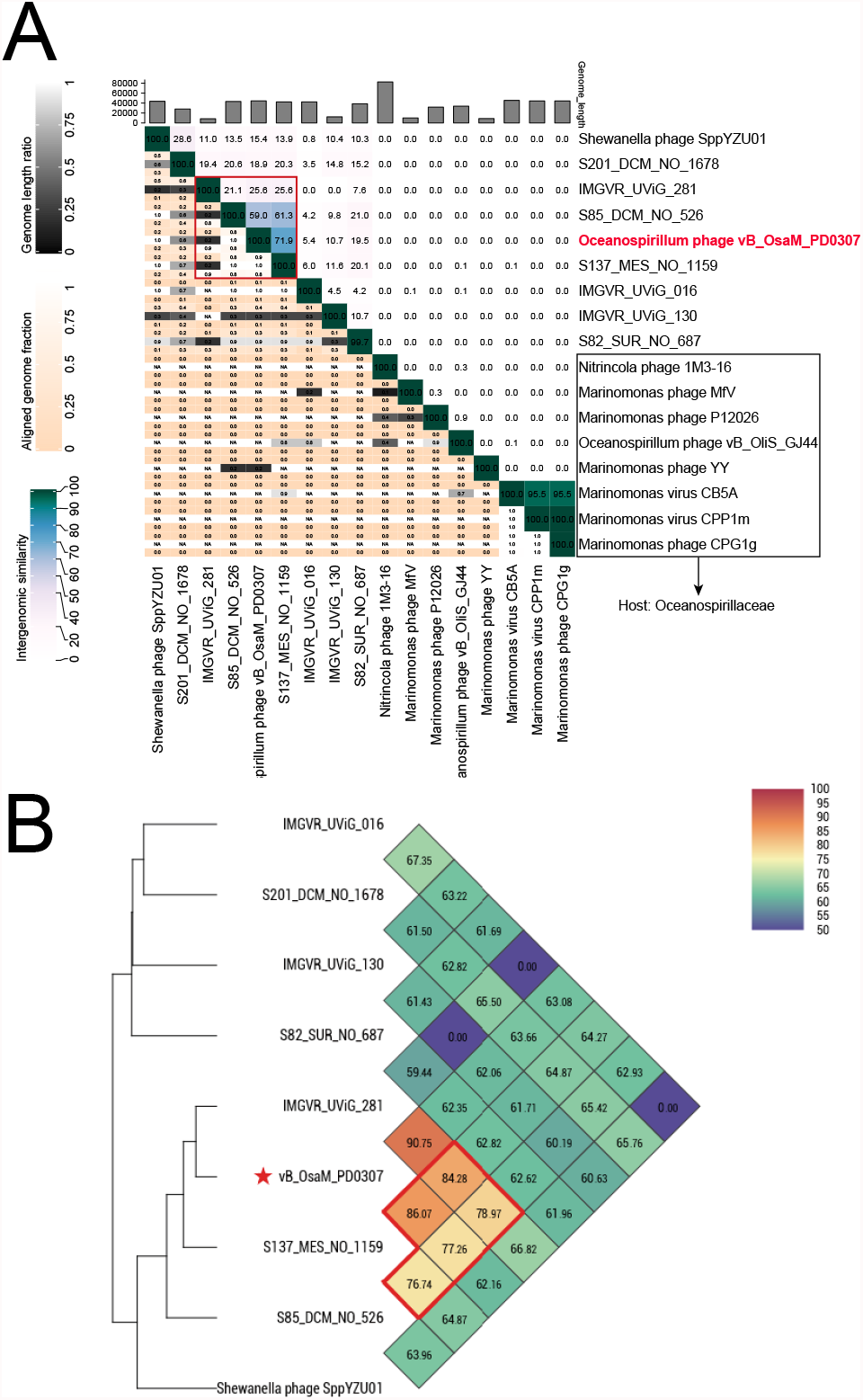
(A) Heatmap of intergenomic similarity values (right half) and alignment indicators (left half and top annotation) of eight *Oceanospirillaceae* phages and eight homologous uncultured viral genomes of Oceanospirillum Phage vB_OsaM_PD0307. The numbers of intergenomic similarity values represent the similarity values for each genome pair, rounded to the first decimal. The genome length ratio for a genome pair using a black to white color gradient indicator. The aligned fraction genome was indicated by orange to white color gradient. (B) The average nucleotide sequence identity (ANI) of Oceanospirillum Phage vB_OsaM_PD0307, Shewanella phage SppYZU01, and seven homologous uncultured viral contigs based on OrthoANI values calculated using OAT software.

To further define the similarity between vB_OsaM_PD0307 and its homologous sequence, the ANI and AAI of all eight homologous sequences were calculated and the results were found to be consistent with VIRIDIC. S85_DCM_NO_526, S137_MES_NO_1159 and IMGVR_UViG_281 were similar to vB_OsaM_PD0307 with high ANI and AAI values (> 70%) (Fig. 6, Table 1, Fig. S1, Table S3). The Bacterial and Archaeal Viruses Subcommittee (BAVS) of the International Committee on the Taxonomy of Viruses (ICTV) considers phages sharing ≥ 70% ANI as members of the same genus (57). As vB_OsaM_PD0307 is the first isolated myovirus infecting *Oceanospirillaceae* and as it is distant from other isolated phages, based on the phylogenetic and synteny analysis, it is suggested that vB_OsaM_PD0307 represents a novel viral genus within the Myoviridae, named *Oceanospimyovirus*. S85_DCM_NO_526 and S137_MES_NO_1159 belong to this new genus together with vB_OsaM_PD0307. IMGVR_UViG_281 might also belong to the genus *Oceanospimyovirus* but having a low-quality genome sequence (Fig. 5B).

**Fig. 6.**
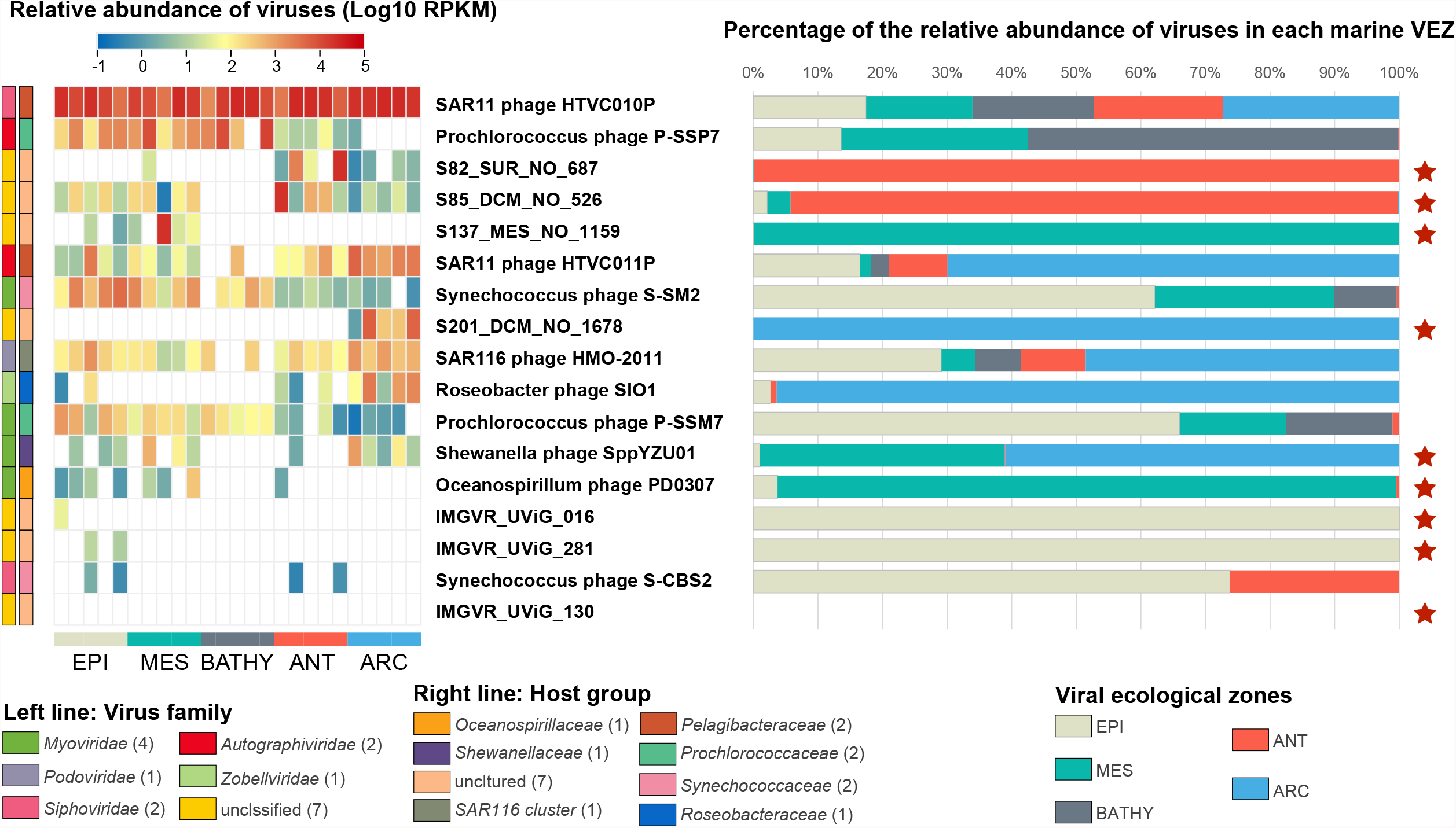
Relative abundance of Oceanospirillum Phage vB_OsaM_PD0307, Shewanella phage SppYZU01, seven homologous uncultured viral contigs and reference phage genomes in the global ocean viromes (GOV 2.0) datasets.

**Table 1.** Amino acid identity of Oceanospirillumphage vB_OsaM_PD0307 and other night homologous sequences.

| Abbreviation | Identity<br>Frequency |  | Bit-score<br>Frequency |  |
| --- | --- | --- | --- | --- |
|  | mean | median | mean | median |
| IMGVR_UViG_3300009431_000281 | 85.08 | 93.7 | 428.9 | 346.5 |
| S137_MES_NO_1159 | 83.36 | 87.01 | 461.6 | 343 |
| S85_DCM_NO_526 | 72.9 | 79.04 | 428.2 | 282 |
| Shewanella phage SppYZU01 | 48.33 | 47.22 | 260.6 | 173 |
| S201_DCM_NO_1678 | 48.24 | 47.73 | 260.4 | 144 |
| IMGVR_UViG_3300003691_000130 | 46.53 | 46.51 | 247.9 | 125 |
| S82_SUR_NO_687 | 45.45 | 44.27 | 249.1 | 159 |
| IMGVR_UViG_3300024520_000016 | 40.97 | 37.32 | 127.7 | 30 |

| Abbreviation | Identity |  | Bit-score |  |
| --- | --- | --- | --- | --- |
|  | Frequency |  | Frequency |  |
|  | mean | median | mean | median |
| IMGVR_UViG_281 | 85.08 | 93.7 | 428.9 | 346.5 |
| S137_MES_NO_1159 | 83.36 | 87.01 | 461.6 | 343 |
| S85_DCM_NO_526 | 72.9 | 79.04 | 428.2 | 282 |
| Shewanella phage SppYZU01 | 48.33 | 47.22 | 260.6 | 173 |
| S201_DCM_NO_1678 | 48.24 | 47.73 | 260.4 | 144 |
| IMGVR_UViG_130 | 46.53 | 46.51 | 247.9 | 125 |
| S82_SUR_NO_687 | 45.45 | 44.27 | 249.1 | 159 |
| IMGVR_UViG_016 | 40.97 | 37.32 | 127.7 | 30 |

### Distribution of vB_OsaM_PD0307 in the global ocean viromes

The biogeographical distribution of vB_OsaM_PD0307 and its closely associated viral sequences was characterized in the Global Ocean Viromes (GOV2.0) data set covering five viral ecological zones (VEZs). The reference genomes include *Shewanella* phage SppYZU01, seven homologous UViGs, and eight typical phages infecting the most abundant bacterial genera, such as *Pelagibacter, Prochlorococcus, Synechococcus, Roseobacter* and S116 cluster. The relative abundances confirmed the high abundances of pelagiphages and cyanophages, as shown in previous studies from Pacific, Indian, and Global Ocean viromes (58–60). vB_OsaM_PD0307 and its associated viral sequences have a wide and diverse distribution.

vB_OsaM_PD0307 and S137_MES_NO_1159 have similar distribution patterns, mostly in MES. Interestingly, S85_DCM_NO_526 and S82_SUR_NO_687 were relatively abundant in the ANT, while S201_DCM_NO_1678 was relatively abundant in the ARC (Fig. 6), which was significantly different from the distribution pattern of vB_OsaM_PD0307. As viral abundance was mostly tightly coupled to that of their host cells, it is proposed that annual marine polar algal blooms might be responsible for the high abundance of *Oceanospirillaceae* (61) and thus the high abundance of *Oceanospimyovirus* in polar oceans (Fig. 7). In coastal waters of the western Antarctic Peninsula, Oceanospirillales and *Pelagibacteraceae* were most abundant in winter and spring. In the Amundsen Sea polynya, *Oceanospirillaceae* is one of the principal dominating bacteria families and in bloom events can contribute up to a 33.9% of the total bacterial 16S rRNA genes (26, 62–64). In the Meade River area of the coastal Arctic, OTUs related to the family *Oceanospirillaceae* comprised the largest component of *Gammaproteobacteria*, approximately 22 and 8% of the bacterial communities in April and August, respectively (29). Although the reasons for the high abundance of host cells in the Arctic and Antarctic are not completely understood, it is hypothesized that: annual algal blooms in polar regions could release large amounts of organic matter that promotes increased bacterial biodiversity (65–67). In addition, algae acquire vitamin B12 through a symbiotic relationship with bacteria (68). Members of *Oceanospirillaceae* have been shown to support phytoplankton growth in polar waters through the synthesis of cobalamin (vitamin B12) (69–72).

Therefore, uncultured viral sequences assembled from the polar viromes might play an important but unrecognized role in regulating the polar bacterial community structure associated with polar algal blooms. This study reinforces the importance and power of the combination of phage isolation and metagenomics to improve our knowledge of marine viral diversity and their ecological significance.

## Conclusion

*Oceanospirillum* has a strong metabolic capacity and a key ecological niche; as such its phage will inevitably affect its abundance, community structure, and metabolic capacity. Here we isolated the first myovirus infecting *Oceanospirillaceae*, named vB_OsaM_PD0307. The presence of TetR family transcriptional regulator encoded by vB_OsaM_PD0307 suggests its potential contribution to the antimicrobial resistance and virulence of its host. vB_OsaM_PD0307 represents a novel myoviral genus-level cluster, named *Oceanospimyovirus*, with two high-quality UViGs. The relative abundance and distribution of *Oceanospimyovirus* suggest that it could be prevalent in polar oceans, coupling with their high-abundant and algal-associated host bacterial communities. The discovery of *Oceanospimyovirus* in the Global Ocean Viromes raised several questions regarding their diversity, ecology, and roles in microbial communities. Here, we performed a culture-based and metagenomics-based analysis of the genomic diversity and distribution of the *Oceanospimyovirus* group. The obtained *Oceanospimyovirus* type genome vB_OsaM_PD0307 helps reveal the genuine extent of the genetic diversity of *Oceanospimyovirus* within natural populations of marine viruses. These novel insights into the diversity and ecology of *Oceanospimyovirus* further expands our current understanding of these important phages. Lastly, further investigation using our newly constructed virus–host models will provide additional valuable insights into the influence of viruses on the interaction among algal blooms, bacteria and viruses in the polar oceans.

## Acknowledgments

We sincerely thank Jia Zhen, School of Computer Science and Technology, Guizhou University, for his help in the data processing. We thank for the support of the high-performance servers of Center for High Performance Computing and System Simulation, Pilot National Laboratory for Marine Science and Technology (Qingdao), the Marine Big Data Center of Institute for Advanced Ocean Study of Ocean University of China, the IEMB-1, a high-performance computing cluster operated by the Institute of Evolution and Marine Biodiversity, and the high-performance servers of Frontiers Science Center for Deep Ocean Multispheres and Earth System.

## Ethical Approval

This article does not contain any studies with animals or human participants performed by any of the authors.

## SUPPLEMENTAL MATERIAL

Supplemental material is available online only.

FIG S1, PDF file, 6321 KB.

TABLE S1, XLSX file, 17 KB.

TABLE S2, XLSX file, 14 KB.

TABLE S3, XLSX file, 28 KB.

**Supplemental flie 1: Fig S1** Amino acid identity and bitscore distribution between vB_OsaM_PD0307, Shewanella phage SppYZU01, and seven homologous uncultured viral genomes

**Supplemental flie 2: Table S1** Genome annotation of Oceanospirillumphage vB_OsaM_PD0307

**Supplemental flie 3: Table S2** The result of BLASTn in IMG/VR and the uncultured contigs‘ information

**Supplemental flie 4: Table S3** Average amino acid identity of Oceanospirillum Phage vB_OsaM_PD0307 between Shewanella phage SppYZU01 and seven homologous uncultured viral genomes

